# *Plasmodium falciparum* PP1 phosphatase is a key regulator of malaria parasite transmission

**DOI:** 10.1101/2025.03.06.641796

**Authors:** Royer Ludivine, Colard-Itté Emma, Tavella Tatyana, Lorthiois Audrey, Goussin Stéphane, N’Dri Marie-Esther, Sabra Reem, Auréline Deiss, Thiberge Sabine, Mauld H. Lamarque, Lavazec Catherine

## Abstract

For the successful transmission of malaria parasites from humans to mosquitoes, *Plasmodium falciparum* gametocytes must remain in the bloodstream long enough to be taken up by a mosquito. Once ingested, they are then activated into gametes to continue the parasite life cycle in the mosquito midgut. Both persistence of gametocytes in the blood and their activation into gametes are tightly regulated by phospho-signaling. While the serine-threonine phosphatase *Pf*PP1 is an essential enzyme for parasite asexual proliferation, its role during transmission of sexual stages remains elusive. Here, we employed a conditional depletion strategy to conduct a functional analysis of *Pf*PP1 during gametocyte development, gamete activation and transmission to mosquitoes. We show that *Pf*PP1 regulates the deformability and the permeability of mature gametocyte-infected erythrocytes through the dephosphorylation of PKA substrates, thus highlighting a key role for *Pf*PP1 in modulating the host cell mechanical properties, which are crucial for gametocyte persistence in the bloodstream. We also provide evidence that *Pf*PP1 controls crucial steps of gamete activation via stimulation of the cGMP/ PKG pathway. Collectively, these results underscore the pivotal role of *Pf*PP1 in the transmission of *P. falciparum* to the mosquito during both sexual development and gamete activation.

**AUTHOR SUMMARY:** The protein phosphatase PP1 is a major contributor to total cellular phosphatase activity in eukaryotes and plays a critical role during various cellular processes. Here, we have unraveled novel mechanisms regulated by the phosphatase *Pf*PP1 in the human malaria parasite *Plasmodium falciparum*. While *Pf*PP1 is known to be essential for the parasite asexual proliferation, in the present study we demonstrate that *Pf*PP1 is also required during the sexual parasite stages, called gametocytes, that ensure parasite transmission from humans to mosquitoes. *Pf*PP1 is involved in regulating the mechanical properties of the gametocyte-infected host cell, a process necessary for the persistence of gametocytes in blood circulation. Moreover, *Pf*PP1 also contributes to the activation of gametocytes into gametes, the stages able to pursue the parasite life cycle in mosquitoes. In addition to providing insights into novel mechanisms involved in parasite transmission, this study also highlights the possibility of interfering with *Pf*PP1 signaling pathway for blocking malarial parasite transmission.

## INTRODUCTION

Malaria, caused by apicomplexan parasites in the genus *Plasmodium*, remains one of the major public health burdens, with more than half a million deaths annually. The clinical symptoms of malaria are attributed to the proliferation of *Plasmodium* asexual stages, whereas parasite transmission from humans to mosquitoes requires that a subpopulation of parasites differentiate into nonreplicating sexual forms termed gametocytes. Reducing the transmission of gametocytes to mosquitoes is now considered crucial for achieving malaria elimination, therefore it is urgent to discover new gametocyte targets for development of drugs to block malaria transmission (1).

For the causative agent of the most severe form of malaria in humans, *Plasmodium falciparum,* gametocyte maturation is divided in five morphologically distinct stages I to V and lasts about 10 days (2). During this period, the morphology of the gametocyte progresses from a round form in stage I to an elongated shape in stages III and IV, finally leading to a crescent shape in the mature stage V. In stages II-III, the elongation of the gametocyte is paired with the elongation of its nucleus, with both processes being regulated by the assembly and disassembly of a network of sub-pellicular and nuclear microtubules (3, 4). Immature gametocytes from stage I to IV sequester in the bone marrow parenchyma and appear only as mature stages in the peripheral blood, where they can persist for several days, available for uptake by mosquitoes (5). The persistence of mature gametocytes in blood circulation is dependent on their ability to decrease the stiffness and the permeability of their host erythrocyte (6–9). Indeed, the newly acquired deformability of gametocyte-infected erythrocyte (GIE) allows mature gametocytes to escape clearance by the spleen, while the virtual absence of channel activity at the erythrocyte membrane likely contributes to reducing the osmotic fragility of mature GIE and may protect them from hemolysis. Then, upon ingestion by mosquitoes, mature gametocytes become rapidly activated and differentiate into gametes. This transformation, called gametogenesis, is activated in the mosquito midgut by the presence of a mosquito metabolite termed xanthurenic acid and/or a rise in pH (10). A decrease in temperature to 20-25;C is essential subsequently to gamete activation (11). Gametogenesis begins with the rapid rounding-up of the crescent-shaped mature gametocytes, is followed by the rupture of the parasitophorous vacuole membrane (PVM) and ends in 15-20 minutes with the egress of male and female gametes from the erythrocyte via an inside-out mode (12). During this process, emerging female macrogametes stay round and undergo minor morphological changes whereas male gametes produce eight flagellated microgametes (13). Male gamete formation is characterized by an atypical rapid mitosis, consisting of three rounds of DNA synthesis and successive spindle formation with clustered kinetochores (14).

Phosphorylation events play a central role during gametocyte development and transmission to mosquitoes since both changes in GIE mechanical properties during gametocytogenesis and activation of gametogenesis are tightly regulated by signaling pathways controlled by kinases and cyclic nucleotides (14–17). For instance, cyclic AMP (cAMP) / protein kinase A (PKA)-dependent signaling has been implicated in the increase of immature GIE stiffness, via PKA-mediated phosphorylation of the parasite proteins STEVOR that interact with the ankyrin complex at the erythrocyte membrane (18–20). Conversely, at maturation, the switch in GIE deformability results from a decrease in cAMP levels due to the high expression of the plasmodial phosphodiesterase *Pf*PDEδ (19, 21). Similarly, cAMP/PKA-dependent signaling is involved in regulating new permeability pathways (NPPs) that are responsible for channel activity at the membrane of erythrocytes infected with gametocytes (6) and asexual stage parasites (22), although the phosphorylated substrates have not been identified yet.

During gametogenesis, increased levels of cyclic GMP (cGMP) activate the cGMP-dependent protein kinase G (PKG) that functions as a master regulator of downstream signaling events leading to PVM rupture (23, 24). Several other kinases involved in regulation pathways controlling gametogenesis have been characterized (14), notably the calcium-dependent protein kinases CDPKs, which are essential for multiple steps of male and female gametogenesis (25, 26), the serine/arginine-rich protein kinase SRPK1, which controls expression of genes crucial for male gamete formation (27), or the mitogen-activated protein kinase MAP-2, which is required for the induction of axoneme motility during exflagellation of male gametes (28).

Unlike kinases, phosphatases have received less attention so far, therefore our knowledge of the role and importance of phosphatases during parasite sexual stage development is still incomplete. Yet, mounting evidence suggests that phosphatases may be involved in many processes in *Plasmodium* sexual stages. Accordingly, a systematic functional analysis of the *Plasmodium berghei* phosphatome revealed that among the 30 predicted protein phosphatases in the genome, 6 are specifically required in sexual stages, thus highlighting the importance of these enzymes for parasite transmission (29). Further functional characterization studies have confirmed that *Pb*PPM1 plays a role in exflagellation, *Pb*PPM5 is crucial for oocyst development, whereas *Pb*PPM2, *Pb*SHLP1 and *Pb*PPKL are implicated in ookinete differentiation and motility (29–33). In addition, among the 16 phosphatases that are essential during asexual blood stages (29), *Pb*PP5 is also required for male gamete fertility, *Pb*PP3 regulates parasite colonization of mosquito midgut cells, while *Pb*PP1 is implicated in the regulation of both mitosis and meiosis in male gamete formation and ookinete differentiation, respectively (32–34). Although most of these studies have been performed in the *P. berghei* mouse model of malaria, several reports also suggest an important role for phosphatases in the sexual stages of the human malaria parasite *P. falciparum*. For instance, treatment of mature GIE with calyculin A, a potent serine/threonine phosphatase inhibitor with high specificity for PP1 and PP2A, increases PKA-dependent phosphorylation of membrane proteins and markedly impairs the deformability of the infected cell (19), suggesting that human or plasmodial PP1 and/or PP2A may contribute to the regulation of GIE mechanical properties during gametocytogenesis. Moreover, *Pf*PP1 is likely to be involved in the mechanisms underlying gamete egress. Indeed, in asexual stages *Pf*PP1 regulates the phosphorylation status of guanylyl cyclase alpha (GCα) and therefore its activity, which is required to stimulate cGMP synthesis and activate its downstream effector PKG (35). This mechanism controls merozoite egress from the red blood cell, a process that shares common pathways with egress of gametes (36). Altogether, these observations compel us to explore whether *Plasmodium* phosphatases, and *Pf*PP1 in particular, could serve as effective targets for transmission-blocking interventions.

In this study, we investigate the potential role of *Pf*PP1 in *P. falciparum* sexual stages. We used the previously described inducible knock-out *pfpp1-iKO* line (35) to carry out a functional analysis of *Pf*PP1 during gametocyte development, gametogenesis and transmission to mosquitoes. We provide evidence that *Pf*PP1 depletion leads to abnormal nucleus organization and dysregulation of GIE mechanical properties throughout gametocyte development, followed by impaired mitosis and host cell egress during gametogenesis. These results highlight that *Pf*PP1 has a crucial function in *P. falciparum* transmission to the mosquito during both sexual development and gamete activation.

## RESULTS

### *Pf*PP1 is the predominant phosphatase expressed in gametocytes

*In silico* analyses and transcriptional studies (https://plasmodb.org) (29, 37, 38) revealed that out of the 28 phosphatases encoded in the *P. falciparum* genome, 12 are more highly transcribed in sexual stages than in asexual stages (Supp. Table 1). Among these 12 phosphatases, *Pf*PP1 is the most highly expressed in stage II and stage V gametocytes. To further characterize the kinetics of *Pf*PP1 expression during the parasite sexual development at the protein level, we conducted a western blot analysis of gametocytes at stages II, III, IV and V with a transgenic line expressing a triple-hemagglutinin (HA_3_) tag at the 3’-end of the endogenous *pfpp1* gene (35). Parasite cultures were treated with N-acetylglucosamine (NAG) to eliminate asexual blood-stage parasites and to obtain synchronous and pure gametocytes. Using anti-HA antibodies, we observed *Pf*PP1 expression at all gametocyte stages, with an increase in expression from stage II to stages IV-V (Supp. Fig. 1A). We next analyzed the subcellular localization of *Pf*PP1 during gametocyte development and gametogenesis. In asexual stages, *Pf*PP1 has been previously localized in both the cytoplasm and the nucleus (35, 39, 40), and in a more indirect manner in the host cell (39). In contrast, our immunofluorescence analysis of PFA-fixed GIE revealed that *Pf*PP1 signal was only observed within the gametocyte but was not detected in the host cell (Fig. 1A). The follow-up of *Pf*PP1 staining during gametocytogenesis revealed a diffuse cytoplasmic pattern of *Pf*PP1 at all gametocyte stages, with a more intense focus near or overlapping the nucleus (Fig. 1B). Upon gamete activation, 2 to 8 distinct foci were observed in association with the dividing nucleus in male gametes, while gametes with only one nucleus exhibited a diffuse *Pf*PP1 staining (Fig. 1C-D and Supp. Fig. 1B). Taken together, these observations suggest that *Pf*PP1 may be associated with the microtubule organizing center (MTOC), the kinetochores and/or other components of the mitotic machinery, which are involved in nucleus elongation during gametocyte development and DNA replication events during male gametogenesis (4, 34).

**Figure 1.**
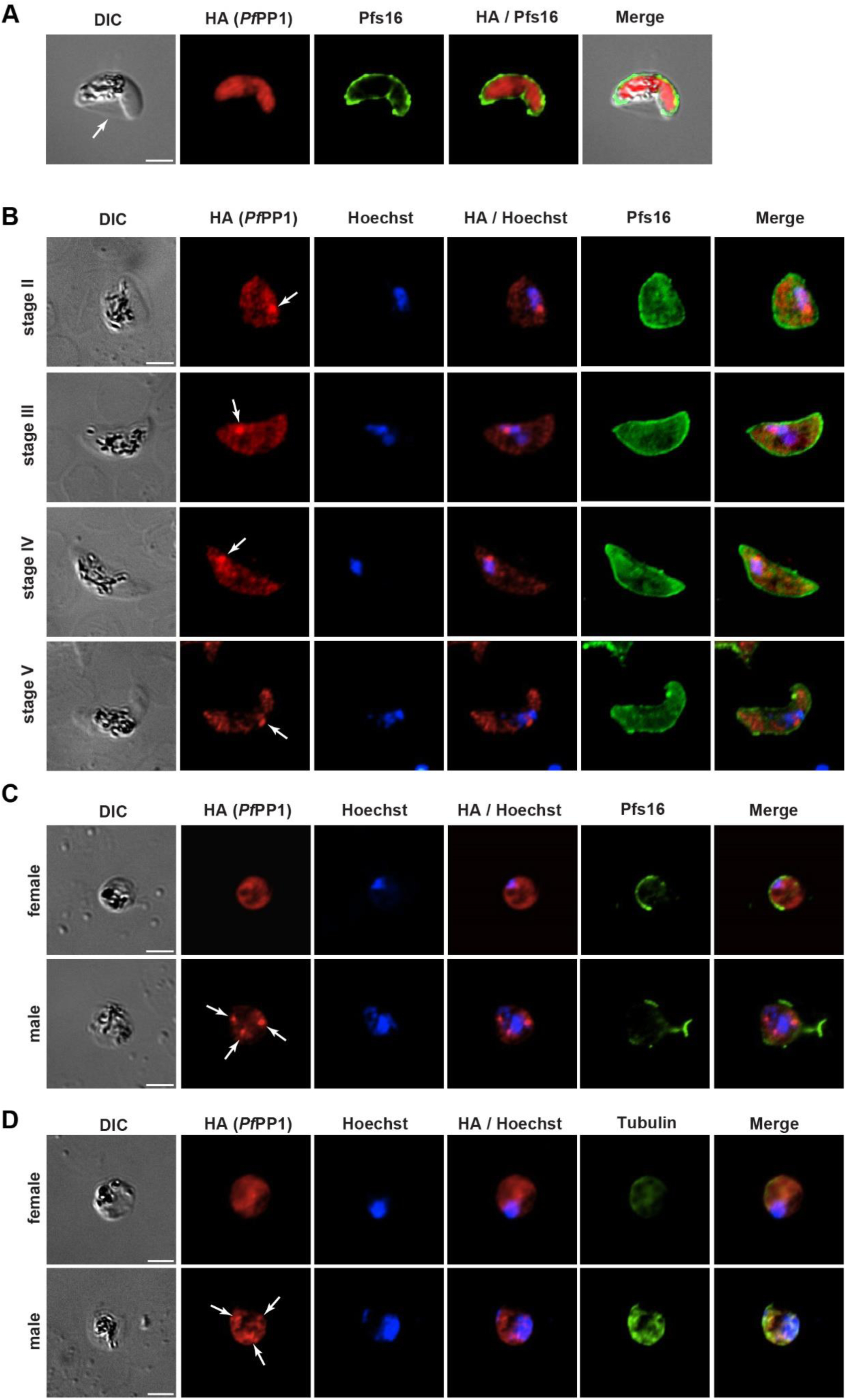
Localization of *Pf*PP1 in gametocytes and gametes. **A.** Immunofluorescence analysis performed on mature (stage V) gametocytes co-labeled with anti-HA antibody and anti-Pfs16 antibody that labels the PVM, showing that *Pf*PP1-HA staining is only detected within the parasite and not in the erythrocyte. The arrow indicates the erythrocyte membrane surrounding the Laveran’s bib. DIC: Differential Interference Contrast. Scale bars: 2µm. **B.** Immunofluorescence analysis performed on stage II, III IV and V gametocytes co-labeled with anti-HA antibody, anti-Pfs16 antibody and Hoechst. The arrows point out *Pf*PP1-HA foci in close proximity to the nuclei. DIC: Differential Interference Contrast. Scale bars: 2µm. **C-D.** Immunofluorescence analysis performed on male and female activated gametes co-labeled with anti-HA antibody, anti-Pfs16 antibody (C), anti-alpha tubulin antibody (D) and Hoechst. Pfs16 staining shows the disappearance of the PVM upon gamete activation. Tubulin staining discriminates male and female gametes. The arrows point out *Pf*PP1-HA foci. DIC: Differential Interference Contrast. Scale bars: 2µm.

### *PfPP1* controls elongation of the nucleus in immature gametocytes

We further carried out a functional analysis of *Pf*PP1 in gametocytes with a conditional depletion strategy. We used the *pfpp1-iKO* line, which is a transgenic line for the inducible knock-out of the *pfpp1* gene, based on the dimerizable Cre recombinase (DiCre) system and expressing a triple-hemagglutinin (HA_3_) tag at the 3’-end of the endogenous *pfpp1* gene, as described by Paul *et al*. (35). We first observed that treatment of *pfpp1-iKO* gametocytes with rapamycin from day 1 post-NAG and over 6 consecutive days induced a complete excision of the *pfpp1* gene from day 3 post-NAG (Fig. 2A), resulting in an 80 % reduction of *Pf*PP1 protein levels at day 6 post-NAG (Fig. 2B). The depletion was also confirmed in mature stages at day 9 post-NAG by immunofluorescence analysis (Fig. 2C). We then investigated whether *Pf*PP1 depletion affects the ability of the parasites to undergo sexual development. First, we did not observe any significative difference in gametocytemia between rapamycin-treated and untreated GIE from day 3 to day 11 post-NAG (Fig. 2D). Furthermore, no differences were identified in the maturation time (Fig. 2E), morphology (Fig. 2F), or sex ratio (Fig. 2G) of *pfpp1-iKO* gametocytes treated or not with rapamycin. Since we observed *Pf*PP1 in association with the nucleus in developing gametocytes (Fig. 1B), we also addressed whether *Pf*PP1 depletion affects the size of the nucleus. Recent ultrastructural analysis of *P. falciparum* gametocytes has described how an intranuclear microtubule network associated with the mitotic machinery leads to the elongation of the gametocyte nucleus, reaching maximum length in stages II/III before being disassembled at the end of stage III, resulting in a more spherical nucleus in stage IV/V (4). Examination of the Hoechst signal in PFA-fixed stage III gametocytes revealed elongated nuclei in DMSO-treated parasites, whereas nuclei exhibited a more spherical profile in rapamycin-treated gametocytes (Fig. 2H). Measurements of the Hoechst signal length in both conditions confirmed that the nucleus size was significantly lower in *Pf*PP1-depleted gametocytes (*p* = 0.0009) (Fig. 2I), suggesting that *Pf*PP1 regulates nuclear elongation during gametocyte development.

**Figure 2.**
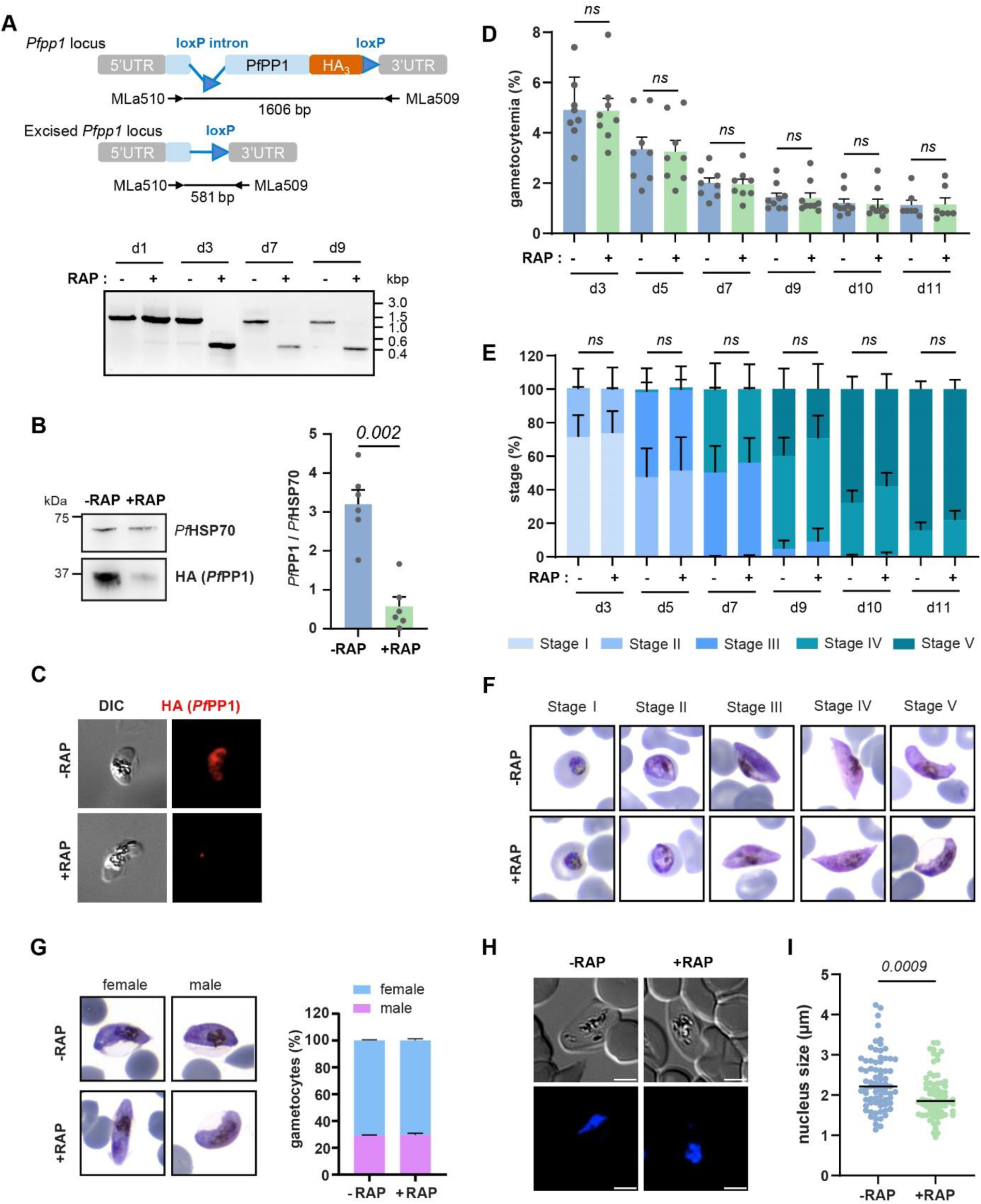
*Pf*PP1 plays no major role in gametocyte maturation but controls elongation of the nucleus in immature gametocytes. **A.** PCR analysis to confirm Rapamycin-induced excision of the *pfpp1* gene in *pfpp1-iKO* transgenic line following rapamycin treatment at day 1, day 3, day 5, day 7 and day 9 post NAG treatment, showing the non-excised (1606 bp) and excised (581 bp) versions of *pfpp1* gene, respectively. Sizes are indicated in kilobase pairs (kbp). **B.** Immunoblot analysis of *Pf*PP1-HA depletion after 6 days treatment with rapamycin (+RAP) or with DMSO (-RAP) in stage III gametocytes using anti-HA antibody. PfHSP70 antibodies were used as control. Bars represent the mean ± SEM from 4 independent experiments (n = 6). Statistical analyses were performed using a Mann-Whitney test. **C.** *Pf*PP1-HA was not detectable by immunofluorescence analysis of stage V gametocytes treated with rapamycin (+RAP) or with DMSO (-RAP). DIC: Differential Interference Contrast. **D-F.** Effect of conditional deletion of *Pf*PP1 on gametocyte development. +RAP and -RAP gametocyte cultures collected on alternate days starting on day 3 post−NAG treatment were scored by microscopy for gametocyte density (D) and distribution of stage I to stage V gametocytes (E). Around 2000 RBCs were scored in each condition. Representative pictures of each gametocyte stages are shown in F. Results show mean ± SEM from 8 independent experiments (n = 8). Statistical analyses for comparison of gametocyte density and distribution of stages were performed using a two-way ANOVA test and using mixed-effects model test, respectively. **G.** The proportion of male and female gametocytes was determined by analysis of Giemsa-stained smears. 100 parasites were scored under each condition. **H-I**. The size of nuclei in stage III gametocytes from the *pfpp1-iKO* transgenic line treated with DMSO (-RAP) or rapamycin (+RAP) was scored by ImageJ using Differential Interference Contrast (DIC) images and Hoechst staining. Each dot represents an individual cell measurement. Statistical analyses were performed using a Mann-Whitney test.

### *Pf*PP1 depletion affects the mechanical properties of mature GIE

During their maturation, *P. falciparum* gametocytes decrease the stiffness and the NPP-dependent permeability of their erythrocyte host through phosphorylation events controlled by PKA (6, 7, 19). Additionally, previous work showed that treatment with Calyculin A, an inhibitor of PP1-and PP2A Ser/Thr phosphatases, can revert the switch in deformability of mature GIE (19, 41, 42), thus suggesting that reversible phosphorylation events may control the mechanical properties of GIE. To address whether reversible phosphorylation is also involved in the decreased channel activity of mature GIE, we investigated the impact of Calyculin A on stage V GIE permeability using isosmotic lysis experiments in sorbitol, a sugar alcohol that permeates through the NPPs (43). After 60 minutes in sorbitol, we observed a drastic increase in the permeability of GIE (*p* = 0.0003) (Fig. 3A), a phenotype that was fully reversed by the addition of the NPP inhibitor 5-nitro-2-(3-phenylpropylamino) benzoic acid (NPPB). These data indicate that the lysis induced by Calyculin A treatment specifically resulted from NPP activation. Since the phosphatase activity targeted by the inhibitor can originate either from the host cell, the parasite, or both, we directly assessed the contribution of *Pf*PP1 to this effect using the *pfpp1-iKO* transgenic line. After 60 minutes in sorbitol, we observed a drastic increase in the permeability of GIE depleted for *Pf*PP1 (*p* < 0.0001) (Fig. 3B), to an extent similar to what was observed with Calyculin A treatment. This phenotype was again fully reverted upon incubation with NPPB. Therefore, these results indicate that the main phosphatase activity responsible for NPP inhibition in mature GIE would correspond to *Pf*PP1 originating from the parasite.

**Figure 3.**
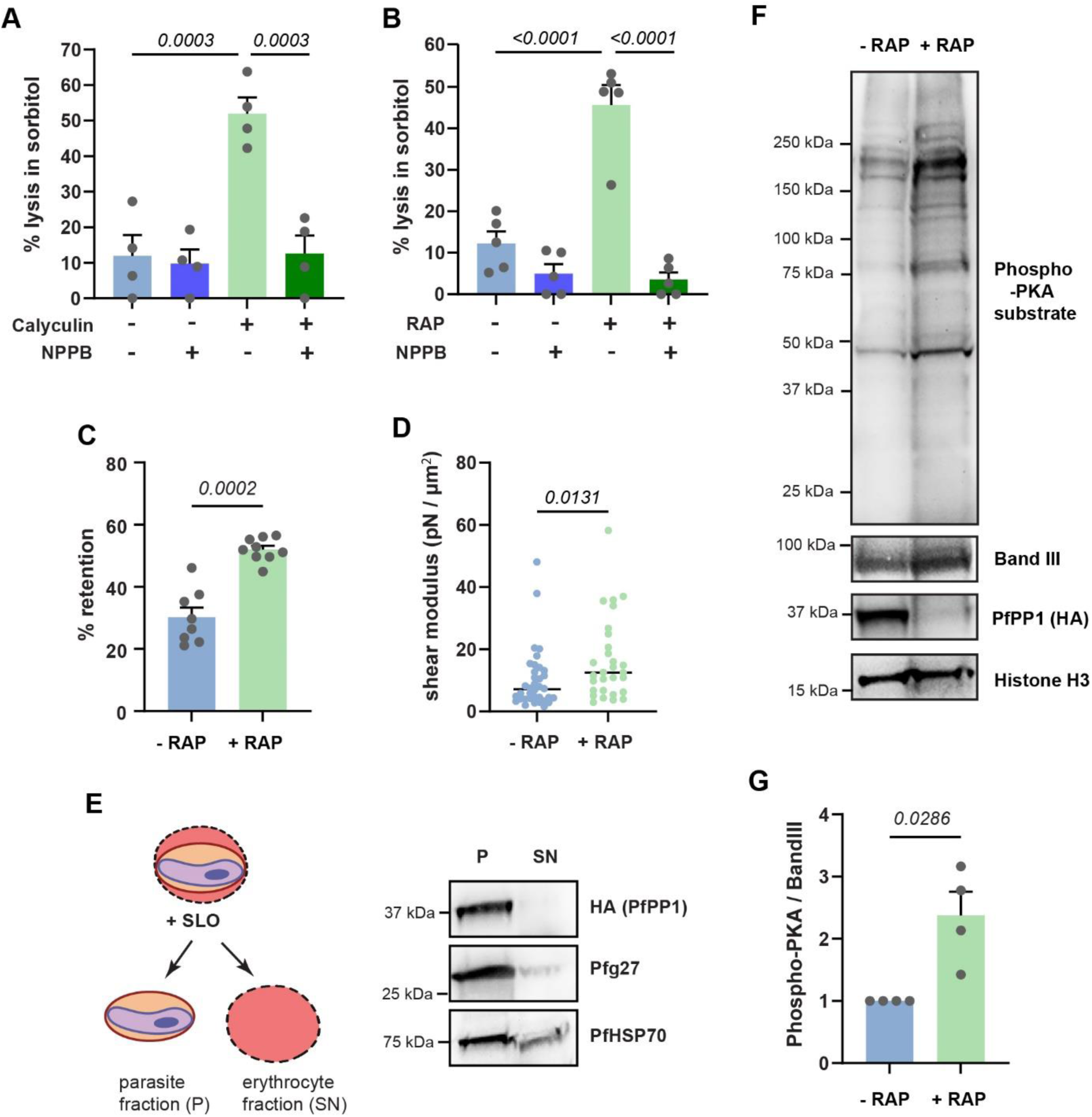
*Pf*PP1 depletion affects the mechanical properties of GIE. **A.** Sorbitol-induced isosmotic lysis of stage GIE from the B10 clone treated for 30 minutes with DMSO or with 50 nM Calyculin A, in the presence or absence of 100 µM NPPB. Bars represent the mean ± SEM from 4 independent experiments (n = 4). Statistical analyses were performed using a one-way ANOVA test. **B.** Sorbitol-induced isosmotic lysis of stages V GIE from the *pfpp1-iKO* transgenic line treated with DMSO (-RAP) or rapamycin (+RAP), in the presence or absence of 100 µM NPPB. Bars represent the mean ± SEM from 5 independent experiments (n = 5). Statistical analyses were performed using a one-way ANOVA test. **C**. Retention in microbeads of stages V GIE from the *pfpp1-iKO* transgenic line treated with DMSO (-RAP) or with rapamycin (+RAP). Bars represent the mean ± SEM from 3 independent experiments in 3 technical triplicates (n = 3). Statistical analyses were performed using a Mann-Whitney test. **D.** Membrane shear moduli measured by micropipette aspiration of stages V GIE from the *pfpp1-iKO* transgenic line treated with DMSO (-RAP) or rapamycin (+RAP). Each dot represents an individual cell measurement. Statistical analyses were performed using a Mann-Whitney test. **E.** Immunoblotting analysis of stage V gametocytes treated with Streptolysin O (SLO). The pellet (P) corresponds to the parasite fraction while the supernatant (SN) corresponds to the erythrocyte fraction, including parasite proteins exported to the erythrocyte. The quality control of parasite and erythrocyte fractions were checked using anti-Pfg27 (middle panel) and anti-*Pf*HSP70 antibodies (lower panel), respectively. The upper panel shows the presence of *Pf*PP1 exclusively in the parasite fraction. **F.** Immunoblot analysis of phosphorylated PKA substrates in MACS-purified stage V GIE from the *pfpp1-iKO* transgenic line treated with DMSO (-RAP) or rapamycin (+RAP). Immunoblot was probed with rabbit monoclonal antibody directed against phospho-PKA substrates (RRX*S/*T), anti-Band 3 antibody to control for red cell quantity, anti-Histone H3 antibody to control for parasite quantity and anti-HA antibody to validate *Pf*PP1 depletion. **G.** Quantification of Phospho-PKA levels relative to Band III, as shown in G, was performed by densitometry (Quantity one software). Statistical analyses were performed using a Mann-Whitney test.

To analyze the role of *Pf*PP1 in the regulation of stage V GIE deformability, we used a matrix of microspheres that mimics the filtration across the spleen (44). In this system, increased retention rates correspond to increased GIE stiffness and impaired filterability. We observed a significant increase in the retention rates of stage V GIE depleted for *Pf*PP1 compared to the control (*p* = 0.0043), indicating that *Pf*PP1 also contributes to the regulation of mature GIE deformability (Fig. 3C). These observations were confirmed with the micropipette aspiration technique, which measures the elasticity of the cell membrane. In this technique, the greater the value of the shear elastic modulus, the more resistant the membrane is to deformation induced by micropipette aspiration (45). The membrane shear elastic modulus of *pfpp1-iKO* stage V GIE treated with rapamycin slightly but significantly increased compared to untreated ones (*p* = 0.013), indicating that the depletion of *Pf*PP1 decreases the elasticity of the erythrocyte membrane in stage V GIE (Fig. 3D). Last, we controlled with the clone B10 (46), derived from the NF54 strain, that the observed changes in mechanical properties did not result from rapamycin treatment (Supp. Fig. 2A-B). Therefore, these results show that *Pf*PP1 regulates both the permeability and the deformability of mature GIE.

### *Pf*PP1 controls mature GIE mechanical properties by modulating PKA-dependent phosphorylation

The impact of *Pf*PP1 on the host cell membrane properties was somewhat surprising given the intracellular localization of the enzyme in gametocytes (Fig. 1A). However, it may result from the export of a fraction of the protein into the host cytoplasm, undetectable by IFA, or alternatively from the regulation of a cAMP/PKA-dependent signalling cascade initiated in the parasite and having repercussions in the host cell. To test the first hypothesis, we selectively permeabilized the erythrocyte membrane with streptolysin O, while leaving the PVM and the parasite plasma membrane intact (47), thus allowing differential extraction of erythrocyte and parasite proteins. Western blot analysis of both fractions revealed that *Pf*PP1 was exclusively found in the parasite fraction (Fig. 3E). Likewise, the parasite cytoplasmic marker Pfg27 (48) was mostly recovered in the parasite fraction, whereas the parasite chaperon protein *Pf*HSP70, which is exported to the erythrocyte (49), was detected in both compartments, as expected. These results do not support a direct role for *Pf*PP1 on GIE membrane properties, although it remains possible that exported protein levels remain below the limit of detection.

Next, we addressed whether the phosphorylation status of PKA substrates, that include host membrane proteins, was altered by *Pf*PP1. We probed extracts of stage V GIE, either deleted for *Pf*PP1 or not, using a monoclonal antibody specific for canonical phospho-(Ser/Thr) PKA substrates sites (RRXS*/T*) (Fig. 3F). The intensity of phosphorylation was 2 to 3-fold higher in *Pf*PP1-depleted parasites, as compared to control, suggesting that *Pf*PP1 may modulate the mechanical properties of mature GIE by interfering with the phosphorylation status of several PKA substrates (Fig. 3G).

Finally, since our previous work showed that decreased stiffness and permeability in mature GIE are correlated with reduced cAMP levels (6, 19) and that *Pf*PP1 depletion lowers cGMP concentration in asexual stages (35), we also investigated whether *Pf*PP1 modulates cyclic nucleotide levels in GIE. To address this hypothesis, we measured cAMP and cGMP levels in *pfpp1-iKO* stage V GIE treated with and without rapamycin. We did not observe any increase of cAMP concentration in mature GIE deleted for *Pf*PP1 (Supp. Fig. 3A), suggesting that the increase in rigidity and the reactivation of NPPs induced by *Pf*PP1 depletion are not mediated by cAMP levels. In addition, cGMP levels were similar with and without rapamycin treatment (Supp. Fig. 3B), suggesting that *Pf*PP1 does not control GIE mechanical properties through modulation of cGMP levels. Consistent with this observation, supplementation with the cGMP analogue 8-bromoguanosine-3’-5’-cyclic monophosphate (8Br-cGMP) did not revert the increase in channel activity induced by *Pf*PP1 depletion (Supp. Fig. 3C). Taken together, these results suggest that *Pf*PP1 modulates GIE mechanical properties through the dephosphorylation of PKA substrates, but not likely through the regulation of cAMP levels or cGMP levels.

### *PfPP1* contributes to parasite transmission to mosquitoes

Based on our results showing that *Pf*PP1-depleted parasites produce stage V GIE, albeit with altered mechanical properties, we wondered whether these mature gametocytes could undergo full development in Anopheles mosquitoes. Mosquitoes were infected by standard membrane feeding assays with NF54 as a control or *pfpp1-iKO* GIE pre-treated with or without rapamycin. Despite similar density of mature gametocytes in reference and *pfpp1-iKO* lines (Fig. 4A), the transmission efficiency of *pfpp1-iKO* GIE was very low compared to NF54 (Fig. 4B and Supp. Table 2). This loss of infectivity could be due to the repeated transfections required to obtain the strain, and precluded any comparison in oocyst density (Fig. 4C). Nonetheless, we observed a significant reduction in oocyst prevalence upon *Pf*PP1 depletion (*p* = 0.0407) (Fig. 4B, Suppl table 2), suggesting that *Pf*PP1 may be important for successful parasite transmission/development in the vector.

**Figure 4.**
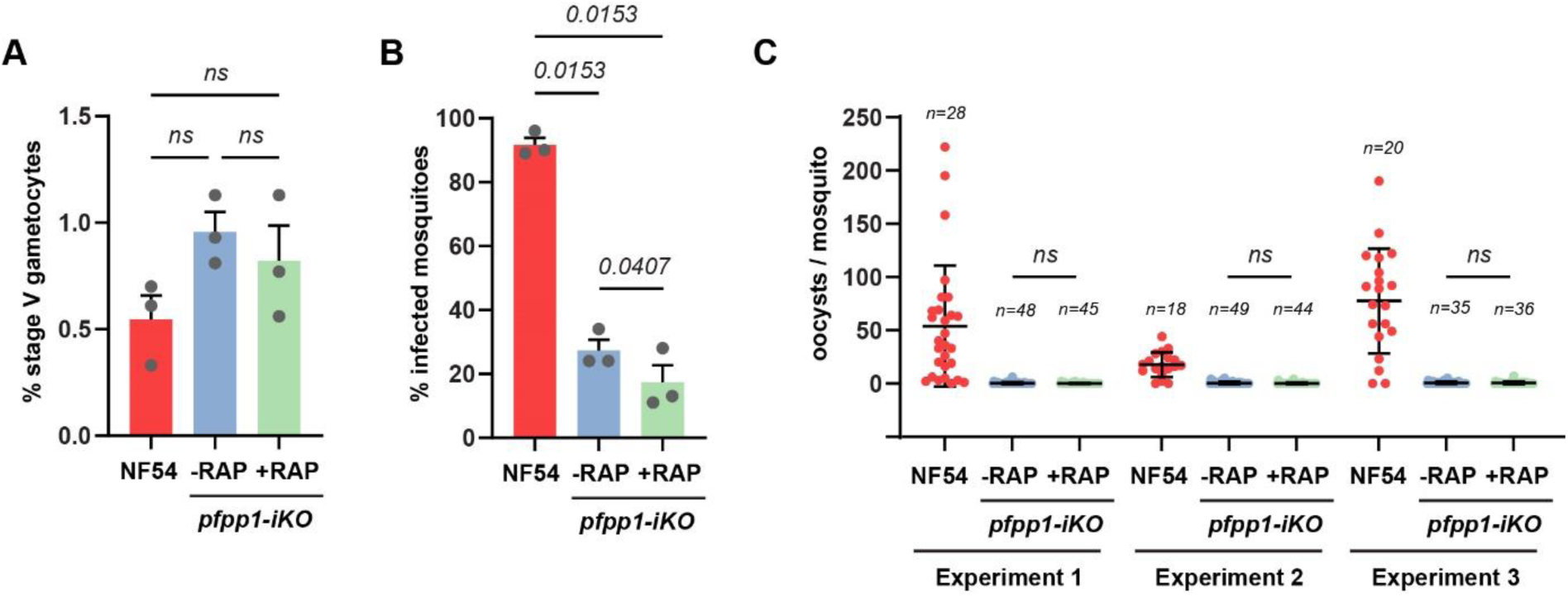
Standard membrane feeding assays of *Anopheles* mosquitoes. **A.** Percentage of stage V gametocytes obtained in cultures from NF54 strain, or *pfpp1-iKO* line treated with DMSO (-RAP) or rapamycin (+RAP), before the mosquito feeding experiments. Bars represent the mean ± SEM from 3 independent experiments (n = 3). Statistical analyses were performed by one-way ANOVA test. **B-C.** *Anopheles stephensi* mosquitoes were fed on cultures of NF54 strain, or *pfpp1-iKO* line treated with DMSO (-RAP) or rapamycin (+RAP). **B.** Mosquitoes were dissected at day 9-11 post-feeding to determine the prevalence of infected mosquitoes. Bars represent the mean ± SEM from 3 independent experiments (n = 3). Statistical analyses were performed by one-way ANOVA test. **C.** The intensity of mosquito infection was scored as the number of oocysts per mosquito midgut at day 9-11 post-feeding. Bars represent the mean ± SD of oocysts par midgut and n represents the number of dissected mosquitoes for each experimental feed. Statistical analyses were performed by one-way ANOVA test.

### *PfPP1* is essential for successful gametogenesis

The decrease observed in oocyst prevalence upon *Pf*PP1 depletion, though moderate, prompted us to investigate gametogenesis, as it might reflect a defect in the capacity to produce fertile gametes. We thus set out to evaluate the effect of *Pf*PP1 depletion at different steps of gametogenesis: DNA replication in male gametes, rounding of gametes, gamete egress from the PV and gamete egress from the erythrocyte. First, to determine whether *Pf*PP1 is involved in mitosis during male gamete formation, as suggested by a recent study conducted in *P. berghei* (34), we monitored DNA replication by immunofluorescence analysis of Hoechst-stained gametes in both rapamycin-treated and untreated parasites (Fig. 5A). The percentage of round gametes that have undergone nuclear divisions at 20 minutes after gamete activation was significantly lower in *Pf*PP1 depleted-parasites than in controls (*p* = 0.0059), suggesting that *Pf*PP1 is involved in the regulation of mitosis in male gametes (Fig. 5B). Next, to score the ability of gametocytes to undergo rounding and egress from the erythrocyte, we used a quantitative gametogenesis assay based on fluorescent wheat germ agglutinin (WGA) surface staining of GIE before exposure to low temperature and increase in human serum concentration (50) (Fig. 5C). This analysis showed that the proportion of rounded gametes 20 minutes after gametogenesis activation was significantly reduced in *pfpp1-iKO* parasites treated with rapamycin compared to untreated ones (*p* = 0.0006) (Fig. 5D). Remarkably, *Pf*PP1 depletion almost totally impaired the capacity of gamete to egress from the erythrocyte (*p* < 0.0001) (Fig. 5E). Gametocytes from the clone B10, treated under the same conditions, did not show any defects in rounding and egress, confirming that the observed effects resulted from *Pf*PP1 depletion and not from rapamycin treatment (Supp. Figure 4A-B). To further address whether gamete egress was impaired before or after PVM rupture, we labeled the PVM using anti-Pfs16 antibodies and quantified the number of Pfs16-negative gametes in the total parasite population after gamete activation (Fig. 5F). Our data revealed that the proportion of PV-egressed gametes drastically decreased upon *Pf*PP1 depletion (*p* < 0.0001), indicating that *PfP*P1 is required for the egress of gametes from the PV (Fig. 5G).

**Figure 5.**
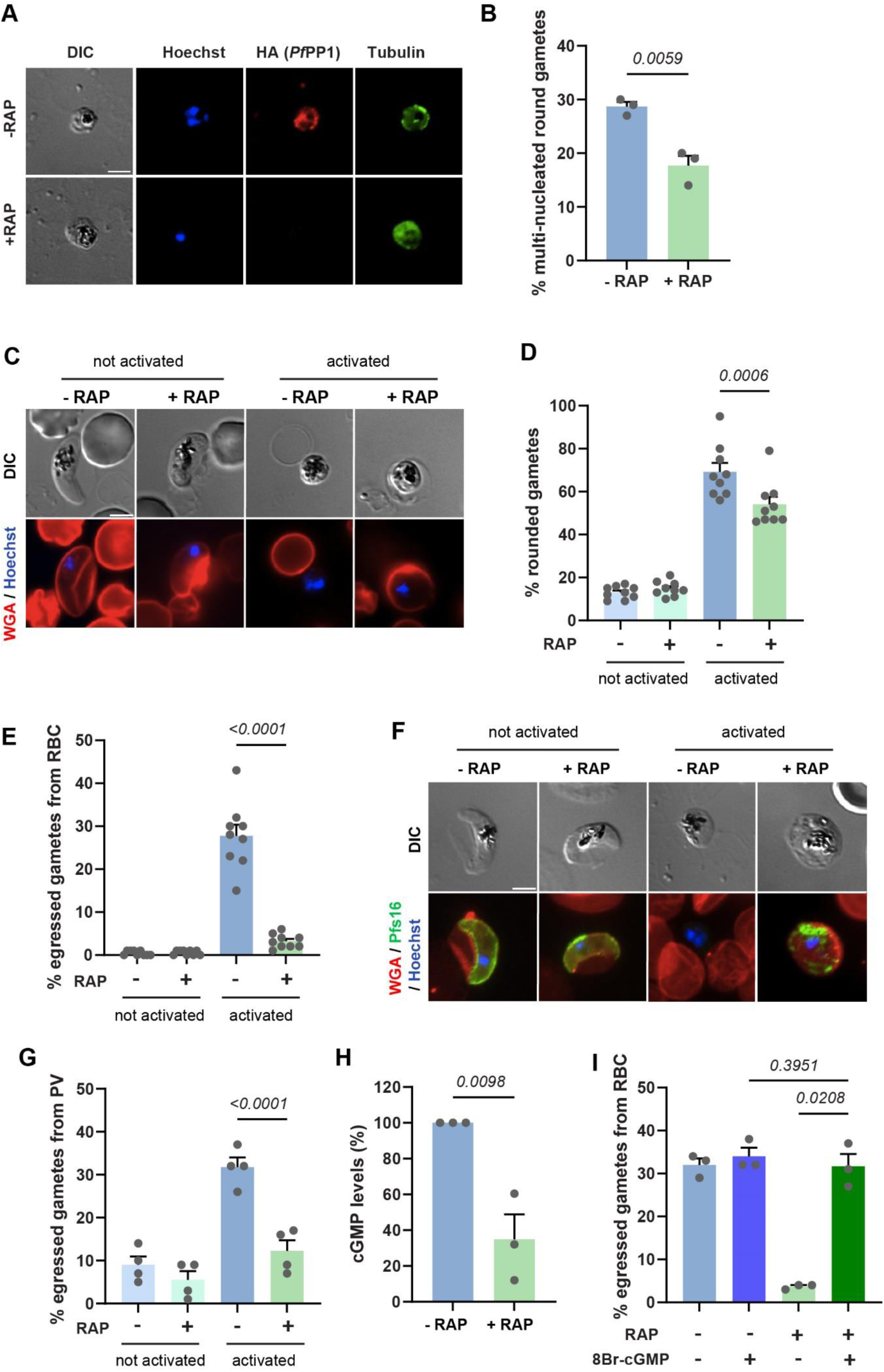
*PfPP1* is essential for successful gametogenesis. **A.** Immunofluorescence analysis of activated gametes from the *pfpp1-iKO* transgenic line treated with DMSO (-RAP) or rapamycin (+RAP). DNA is stained with Hoechst. Anti-HA staining confirms *Pf*PP1 depletion and tubulin staining shows male gametes. Scale bar: 2 µm. **B.** Quantification of nuclear divisions 20 minutes after gamete activation shown as the percentage of round cells with dividing nucleus in parasites treated (+RAP) or not (-RAP) with rapamycin. Bars represent the mean ± SEM from 3 independent experiments (n = 3). Statistical analyses were performed by unpaired t-test. **C-E.** *pfpp1-iKO* gametes rounding and egress from the erythrocyte were scored using Differential Interference Contrast (DIC) images and WGA-Texas Red staining to detect the erythrocyte membrane. Stage V GIE treated with DMSO (-RAP) or rapamycin (+RAP) were sampled before (not activated) and 20 min following activation (activated). Red: WGA-Texas Red, blue: Hoechst. Scale bar: 2 µm. At least 100 parasites were scored under each condition to determine the percentage of rounding up (D) or egress from the erythrocyte (E). Bars represent the mean ± SEM from 9 independent experiments (n = 9). Statistical analyses were performed by one-way ANOVA test. **F.** Gamete egress from the PV was scored using Pfs16 antibodies to detect the PVM. Red: WGA-Texas Red, green: Pfs16, blue: Hoechst, DIC: Differential Interference Contrast. Scale bar: 2 µm. **G.** Percentage of egressed gametes from the PVM before and 20 minutes post-activation of stage V GIE from the *pfpp1-iKO* transgenic line treated with DMSO (-RAP) or rapamycin (+RAP). Bars represent the mean ± SEM from 4 independent experiments in which 100 parasites were scored under each condition (n = 4). Statistical analyses were performed by one-way ANOVA test. **H.** Relative cGMP levels 20 min after activation in gametes from the *pfpp1-iKO* transgenic line treated with DMSO (-RAP) or rapamycin (+RAP). Bars represent the mean ± SEM from 3 independent experiments (n = 3). Statistical analyses were performed by unpaired t-test. **I.** Percentage of egressed gametes from the erythrocyte before and after activation of stage V GIE from the *pfpp1-iKO* transgenic line treated with DMSO (-RAP) or rapamycin (+RAP), supplemented or not with 1 µM 8Br-cGMP. Bars represent the mean ± SEM from 3 independent experiments (n = 3). Statistical analyses were performed by one-way ANOVA test.

Interestingly, the defects in gamete rounding and egress induced by *Pf*PP1 depletion are reminiscent of the phenotype observed upon inhibition of PKG (23). Since *Pf*PP1 has been shown to regulate egress of asexual stages by stimulating GCα activity (35), we measured the effect of *Pf*PP1 depletion on cellular cGMP levels after gamete activation. We found that the depletion of the phosphatase enzyme reduces the second messenger in activated gametes by ∼3-fold (Fig. 5H). Accordingly, supplementation with the cGMP analogue 8Br-cGMP totally reverted the inhibition of gamete egress in *pfpp1-iKO* cultures (Fig. 5I), suggesting that *Pf*PP1 stimulates gamete egress by controlling cGMP levels. Collectively, these results show that *Pf*PP1 governs the gamete activation pathway leading to egress from the host cell, likely via stimulation of the cGMP/ PKG pathway.

## DISCUSSION

Phospho-signaling plays crucial roles in the development of *P. falciparum* gametocytes and their activation into gametes, both of which are key steps in the successful transmission of the parasite from humans to mosquitoes. The *P. falciparum* genome encodes for a set of protein kinases and phosphatases that regulate various processes throughout different stages of the parasite life cycle (51). Among these, *Pf*PP1 is the most quantitatively significant protein phosphatase expressed in sexual stages (29, 37, 38). In this study, we functionally characterized *Pf*PP1 during gametocytogenesis, gametogenesis and transmission to mosquitoes. Our results revealed that *Pf*PP1 regulates both the elongation of the gametocyte nucleus and the mechanical properties of the infected host cell during gametocyte development. Additionally, it contributes to multiple steps of gametogenesis, including rounding of gametes, gamete egress from the PV, gamete egress from the erythrocyte and DNA replication in male gametes. Interestingly, our data suggest that *Pf*PP1 contributes to these processes through distinct pathways. Specifically, it regulates GIE mechanical properties via the cAMP/PKA pathway, while it controls the initiation of gametogenesis by activating the cGMP/PKG pathway.

The observed increase in phosphorylation of PKA substrates in *Pf*PP1-depleted GIE suggests that *Pf*PP1 may contribute to the switch in deformability and to the decrease of NPP activity in mature GIE by modulating PKA substrate phosphorylation. However, it remains unclear whether the proteins dephosphorylated by *Pf*PP1 are directly involved in regulating GIE mechanical properties and whether they originate from the erythrocyte or from the parasite. Although *Pf*PP1 depletion results in a decreased elasticity of the erythrocyte membrane, our subcellular localization studies indicate that the enzyme is not exported by gametocytes into the host cell, making it unlikely that *Pf*PP1 directly dephosphorylates erythrocyte proteins or parasite proteins located at the erythrocyte membrane. Instead, we favor the hypothesis that *Pf*PP1 may activate a signaling cascade, eventually leading to the dephosphorylation of proteins involved in regulating the mechanical properties at the erythrocyte membrane. The final substrates of this signaling cascade may be the parasite proteins STEVOR, which have been shown to be phosphorylated by PKA, to interact with the erythrocyte cytoskeletal ankyrin complex and to regulate GIE deformability (20). Additionally, we can also hypothesize that *Pf*PP1 activity leads to the dephosphorylation of erythrocyte cytoskeleton proteins crucial for the mechanical properties of infected erythrocytes. In support of this hypothesis, analysis of the phospho-proteome of erythrocytes infected by parasites expressing a conditional destabilized version of *Pf*PP1 at the schizont stage revealed a significant increase in phosphorylation of several erythrocyte cytoskeletal proteins, including adducin (35). Notably, adducin is a known substrate of PKA (52) and promotes the binding of spectrin to actin, which regulates erythrocyte membrane properties (53). Further work is needed to determine whether adducin phosphorylation directly contributes to the loss of deformability in *Pf*PP1-depleted GIE.

A recent study proposed that microtubule-based deformations of the nucleus could play a role in the rigidification of immature gametocytes (4). However, it seems unlikely that *Pf*PP1 regulates GIE mechanical properties through this mechanism since *Pf*PP1-depleted GIE display increased stiffness alongside a shorter nucleus. Interestingly, the same study also proposed that nuclear elongation may play a role in controlling gene transcription, as previously shown in hematopoietic stem and progenitor cells (4, 54). While this hypothesis requires further investigation, it offers an exciting possibility, positioning *Pf*PP1 as a potential master regulator of gene expression in developing gametocytes.

Gametogenesis is a complex, multi-step mechanism that is finely regulated. We observed that *Pf*PP1 depletion interferes with different steps of gamete activation, including rounding-up, egress from the PV and the erythrocyte, and nuclear divisions in male gametes. The phosphatase PP1 is known to be a pleiotropic enzyme in mammals (55) and in *Plasmodium* (51), so it is plausible that *Pf*PP1-mediated dephosphorylation regulates gametogenesis through multiple mechanisms. Consistent with this hypothesis, we recently reported that *Pf*PP1 is involved in two distinct steps of the egress pathway in asexual stages: the initiation of the rounding-up of the PV through calcium signaling and the rupture of the PVM through modulation of cGMP levels (56). Moreover, in addition to its role in host cell egress, *Pf*PP1 is also required for schizogony during the schizont stages (35). We hypothesize that *Pf*PP1 may regulate mitotic divisions during male gametogony by phospho-regulation of motor proteins that play an important role in spindle assembly and axoneme formation, as previously suggested in *P. berghei* (34, 57), whereas it may contribute to the activation of gametogenesis by phospho-regulation of GCα, as shown in asexual stages (35). Mounting evidence suggests that the same membrane signaling platform is at play to regulate GC activity in different parasite stages across the phylum of Apicomplexa (16). Specifically, several studies revealed that GCα activity depends on several co-factors, including GEP1, UGO, SLF and CDC50B (11, 58–60). Although it is still unknown whether *Pf*PP1 is part of this signaling platform, future work should investigate whether *Pf*PP1 regulates the interactions between GCα and its co-factors, and whether GCα-interacting partners are also substrates for *Pf*PP1.

Mosquito infection experiments showed that *Pf*PP1 depletion resulted in a modest yet significant reduction in parasite development within mosquitoes. However, this outcome is challenging to interpret because of the exceptionally low transmission efficiency of *the pfpp1-iKO* parasite line, which may have been compromised during transfection experiments. Nonetheless, despite the fact that mosquito infection experiments were not fully conclusive, the strong inhibition of gamete egress and the defect in mitosis in male gametes caused by *Pf*PP1 depletion opens avenues for developing drugs that target *PfP*P1 activity to block malaria parasite transmission to mosquitoes. Additionally, the increased retention of *Pf*PP1-depleted mature GIE in a spleen mimetic filtration system, which has been validated in human spleens (61) and in humanized mice (62), suggests that inhibitors of *Pf*PP1 may eliminate mature gametocytes from the peripheral circulation, thus making gametocytes unavailable for blood-feeding mosquitoes. This strategy has been previously proposed for other parasite targets (41, 42, 62, 63). Moreover, our previous work suggested that the reactivation of channel activity in mature GIE may promote their hemolysis, which may be another way to eliminate gametocytes from the circulating blood (6, 21). In summary, our results suggest that drugs targeting *Pf*PP1 activity could block parasite transmission by interfering with multiple mechanisms. Since PP1 is highly conserved across organisms, including *Plasmodium* (51), designing *Plasmodium*-specific drugs targeting *Pf*PP1 catalytic domain presents challenges. However, several studies pointed out that *Pf*PP1 forms complexes with hundreds of regulators, which provide a specific location and a selective function for this enzyme (64), as demonstrated for the Gametocyte EXported Protein 15 (GEXP15) (65). Therefore, developing small molecules that interfere with the ability of *Pf*PP1 to bind to its regulators could offer a promising strategy for designing new antimalarials aimed at blocking malaria transmission to mosquitoes.

## MATERIELS AND METHODS

### Parasite culture and gametocyte production

*P. falciparum* parasites from the *pfpp1*-iKO line (35) and from the B10 clone (46) were cultivated *in vitro* under standard conditions using RPMI 1640 medium supplemented with 10% heat-inactivated human serum and human erythrocytes at a 5% hematocrit. Parasites were kept at 37°C in a 5 % O_2_, 5 % CO_2_ and 90 % N_2_ atmosphere. Synchronous production of gametocytes was achieved by increasing parasitemia up to 10-20 % and treating synchronized cultures at the ring stage (day 0) with 50 mM N-acetylglucosamine (NAG) to eliminate asexual parasites (66). NAG was maintained for 6 days until no asexual parasites were detected in the culture. Giemsa-stained smears were scored daily for gametocyte density and distribution of gametocyte stages (stages I to V).

### Immunoblot analysis

Synchronized gametocytes from each stage were purified either by a Percoll gradient composed of 63% of Percoll (Sigma Aldrich St. Louis, MO, USA), 30% of RPMI 1640 medium and 7% of PBS 10X or by magnetic isolation using a MACS depletion column (Miltenyi Biotec) in conjunction with a magnetic separator. 5x10^6^ purified gametocytes were denatured in lysis buffer (XT Sample Buffer, Biorad) in the presence or absence of reducing agent (Biorad), heated for 5 min at 90°C and then separated on 4-12 % gradient polyacrylamide gels (Biorad). After 1 h 30 migration at 150 V, proteins were transferred to PVDF membranes (Biorad) for 1 h at 100 V. After blocking for 2-3 h in a solution of PBS 1X / Tween 20 0.005 % / 5 % skimmed milk powder (Régilait), antibody reactions were carried out in in PBS 1X / Tween 20 0.005 % / 5 % skimmed milk and washing in PBS 1 X / Tween 20 0.005 %. Immunoblots were probed overnight with one of the following primary antibodies: anti-HA rat mAb (clone 3F10, Roche) at 1:1000, anti-Histone H3 rabbit polyclonal antibody (abcam) at 1:500, anti-Band3 mouse mAb (clone A-6, SantaCruz) at 1:1000 or anti-Phospho-PKA substrate (RRXS*/T*) rabbit mAb (clone 100G7, Cell Signaling Technology) at 1:1000. After 3 washed in PBS 1X / Tween 20 0.005 %, immunoblots were incubated 1 hour with horseradish peroxidase-conjugated anti-rat, anti-rabbit or anti-mouse IgG secondary antibodies (Promega) at 1:10,000. Bands of interest were visualized by chemiluminescence after the addition SuperSignal™ West Pico PLUS Chemiluminescent Substrate (Thermo Fisher). The intensity of the bands was quantified with ImageJ, Quantity One or Multigauge software.

### Subcellular fractionation

To determine the subcellular localization of PfPP1, 5x10^6^ MACS-purified mature gametocytes from the *pfpp1*-iKO line were treated for 30 min at 37°C with 150 Units of Streptolysin O. Streptolysin O was activated using 10 mM DTT in 1X PBS before addition to infected erythrocytes. Cells were pelleted at 1,500 *g*, washed once in PBS and resuspended in a volume equivalent to the supernatant fraction. Equal amounts of pellet (parasite fraction) and supernatant (erythrocyte fraction) were analyzed by immunoblotting using anti-HA rat mAb (clone 3F10, Roche) at 1:1000, anti-Pfg27 rabbit serum (48) at 1:1000 and anti-PfHSP70 mouse mAb (20)at 1:1000.

### Immunofluorescence assays

200 µl culture were fixed with 4 % paraformaldehyde / 0.0075% glutaraldehyde in 1X PBS for 15 min at room temperature. After one wash in 1X PBS, cells were permeabilized with 0,1 % Triton X100 for 30 min at 4°C. Cells were then washed again in 1X PBS, spotted on 10-well slide and dried. Alternatively, blood smears were air-dried, fixed with 1 % paraformaldehyde and permeabilized with 0,1 % Triton X100 for 30 min at 4°C. The non-specific binding sites were saturated for 1 h in 1X PBS / 2 % BSA. The smears were then incubated overnight with anti-HA rat mAb at 1:200, anti-Pfs16 rabbit serum (BEI resources) at 1:500 or anti-alpha-tubulin mouse antibody (Sigma) at 1:500 in 1X PBS / 2 % BSA for 2 h at room temperature. After three washes in 1X PBS the smears were incubated with Alexa594 anti-rat and Alexa488 anti-rabbit secondary antibodies at 1:2000 (Life Technologies) and DNA intercalating dye Hoechst 33342, at 1:20000 (Life technologies). Smears were observed at 100X magnification using a Leica DMi8 fluorescence microscope. The sizes of gametocyte nuclei were measured by analyzing Hoechst-stained slides with the ImageJ software.

### Conditional depletion of PfPP1

Synchronized stage I gametocytes of the *pfpp1*-iKO line were treated with 10 nM rapamycin or 0.04% DMSO for 5 days (from day 1 to day 6 post NAG). Gene excision was validated by PCR at day 3, day 5, day 7 and day 9 post NAG with the following primers: MLa510 5’ catggacagttttatgatttgttaagg 3’ and MLa509 5’ gaaaaatactacttttatagataatatttgttttgttc 3’. Protein depletion was validated at day 6 post NAG by immunoblot and at day 9 post NAG by immunofluorescence analysis using anti-HA antibodies.

### Microsphiltration

To address the role of PfPP1 on deformability of gametocyte-infected erythrocytes, parasite cultures containing 1-3 % stage V were subjected to microsphiltration experiments on tips(44). In brief, 400 µl of a mixture of tin beads ranging in size from 5 to 25 µm (Industrie des Poudres Sphériques) were deposited on the filter of a cone acting as a support. Suspensions of cultures containing 1-3 % of Stage V gametocytes were perfused through the microsphere matrix at a flow rate of 60 mL/h using an electric pump (Syramed_sp6000, Arcomed_Ag), followed by a wash with 5 mL of complete medium. The upstream and downstream samples were collected and smeared onto glass slides for staining with Giemsa reagent, and parasitemia was assayed by counting 2000 erythrocytes. Retention on the beads, which is inversely proportional to the deformability of the cells, was calculated by determining the ratio of the parasitemia after bead passage to the parasitemia before bead passage.

### Micropipette aspiration

To address infected erythrocyte membrane elasticity, parasite cultures containing stage V gametocytes were washed and resuspended in pre-warmed 1X PBS / 0.4 % BSA, and then set into a sample chamber placed on the stage of an inverted light microscope (Nikon Diaphot). A glass micropipette mounted on a hydraulic micromanipulator (Narishige, Japan) was used to extract the GIE membrane shear modulus by micropipette aspiration technique. The membrane shear elastic modulus (units: pN/µm) was accessed by the formula:

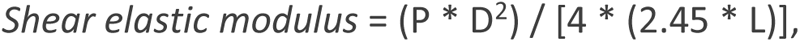

where L represents the length of the membrane tongue aspirated, D corresponds to the micropipette diameter, and P corresponding to the applied pressure in Pa (67). The cell membrane was monitored using a 40X lens. Images were captured using a Nikon D5600 Camera (Nikon). The images were analyzed with ImageJ software for obtention of L and D measurements.

### Isosmotic lysis assays

To test sorbitol sensitivity, 500 µl of a culture containing 1-3 % stage V gametocytes were washed with RPMI and resuspended in 500 µl isosmotic solution containing 300 mM Sorbitol, 10 mM HEPES, 5 mM glucose, with a pH adjusted at 7.4, with or without the addition of 100 µM of the New Permeability Pathways inhibitor 5-nitro-2-(3-phenylpropylamino) benzoic acid (NPPB)(68). After 60 min, samples were smeared onto glass slides for staining with Giemsa reagent, and parasitemia was assayed by counting 2000 erythrocytes. The percentage of gametocyte lysis was obtained by calculating the ratio of parasitemia before and after incubation in the sorbitol solution.

### Gametogenesis assays

At day 10-11 post NAG, stage V gametocytes from the *pfpp1*-iKO line treated with rapamycin or DMSO were subjected to gametogenesis assays (50) with slight modifications to quantity parasite rounding, nuclear divisions and parasite egress from the host cell and from the PVM. For quantitative assessment of parasite rounding and parasite egress from the erythrocyte host, 200 µl samples from each condition were stained for 15 minutes at 37°C with 5 µg/ml Wheat Germ Agglutinin (WGA)-Alexa Fluor 594 (Invitrogen) and 0.5 µg/ml Hoechst 33342. Samples were then pelleted at 1,000 g for 1 min, then mixed in 50 µl human serum and subjected to a temperature drop (from 37 °C to 22 °C / room temperature) for 20 minutes during which gametogenesis took place. Samples were fixed either before or 20 min after gametogenesis activation with 1 % paraformaldehyde for 30 min at room temperature. After one wash in 1X PBS, stained cells were either analyzed with a Fortessa cytometer (Becton Dickinson®) using the FlowJo® program for data analysis to quantify nuclear divisions, or were mounted in a microscope slide under a sealed coverslip. Percent rounding and gamete egress were determined by calculating the % of round gametes and of WGA-negative gametes (due to loss of host erythrocyte) in the total gametocyte population, respectively. For quantitative assessment of parasite egress from the PVM, samples were fixed either before or 20 min after gametogenesis activation with 4 % paraformaldehyde / 0.0075% glutaraldehyde in PBS for 15 min at room temperature, then washed once in 1X PBS and permeabilized with 0,1 % Triton X100 for 30 min at 4°C. Cells were then washed again in 1X PBS, spotted on 10-well slide, dried, and analyzed by immunofluorescence assay using anti-Pfs16 rabbit serum. Percent gamete egress from the PVM was determined by calculating the % of Pfs16-negative gametes in the total gametocyte population. For all the samples, at least 100 gametocytes were analyzed.

### Measurement of cGMP and cAMP levels

Intracellular cGMP levels in mature gametocytes and gametes were measured using enzyme-linked immunosorbent assay (ELISA)-based high-sensitivity direct cGMP/cAMP colorimetric assay kits (Enzo Life Sciences). Gametocytes were obtained through magnetic purification and percoll-purification of rapamycin- and DMSO-treated cultures, 11 days after NAG treatment. Half of the gametocytes were activated with 50µL of human serum for 20min to promote egress and then washed with PBS. Parasites were then pelleted at 9,000 x g, resuspended in 500 µl of 0.1 M HCl, and incubated at room temperature for 10 minutes with intermittent vortexing to ensure complete cell lysis. The samples were pelleted at 9,000 x g, and the supernatant was collected snap-frozen in liquid nitrogen, and stored at –80°C until further use. Samples and standards were acetylated in order to enhance sensitivity according to the manufacturer’s instructions. For each sample, 1.25 × 10⁶ parasites were used. The detection range was 0.08 to 50 pmol/mL for the cGMP assays and 0.078 to 20 pmol/mL for the cAMP assays. All samples and standards were set up in duplicate. Absorbance was measured at 405 nm using TECAN Infinite M1000 Pro plate reader. Sample cyclic nucleotide levels were estimated using logit-transformed linear fits of the measurements obtained from the provided cGMP/cAMP standards.

### Mosquito infections

Standard Membrane Feeding Assays (SMFA) were performed to determine the impact of *Pf*PP1 depletion on gametocyte infectivity for mosquitoes. At day 10-11 post NAG, stage V gametocytes from the *pfpp1*-iKO line treated with rapamycin or DMSO or from the NF54 strain were resuspended in human serum AB+ and fresh erythrocytes (vol/vol) and fed, using a membrane feeder, to 2-to 4-d-old female *Anopheles stephensi*. Mosquitoes were kept at 26°C, 70% humidity, and 10% sucrose. Mosquito infection prevalence (number of infected mosquitoes) and intensity (number of parasites per mosquito) were determined by counting oocysts in mosquito midguts stained with 0.25% fluorescein on day 9-11 post-feeding.

### Statistical analyses

Statistical analyses were performed using GraphPad Prism Version 10.2.3 for Windows.

## ACKNOWLEDGEMENTS

This work was supported by the Agence Nationale de la Recherche (ANR-21-CE15-0043), and the Laboratoire d’Excellence Parafrap (ANR-11-LABX-0024). The authors also thank the financial support of the Centre National de la Recherche Scientifique, the Institut National de la Santé et de la Recherche Médicale, the Université Paris Cité and the Fondation pour la Recherche Médicale. We thank Jérome Clain for kindly providing rapamycin.

## AUTHOR CONTRIBUTION

C.L. and M.H.L. conceived the project and acquired the funding. L.R., E.C.I., T.T., A.L., S.G., M.E.N., R.S. and A.D. performed the experiments. C.L., L.R. and E.C.I designed and interpreted the experiments. M.H.L. and S.T. contributed resources or data. C.L. and L.R. wrote the article, with major input from M.H.L.

